# Sensorimotor cortical-subthalamic network dynamics during force generation

**DOI:** 10.1101/306332

**Authors:** Ahmad Alhourani, Anna Korzeniewska, Thomas A. Wozny, Witold J. Lipski, Efstathios D. Kondylis, Avniel S. Ghuman, Nathan E. Crone, Donald J. Crammond, Robert S. Turner, R. Mark Richardson

## Abstract

The subthalamic nucleus (STN) is proposed to participate in pausing, or alternately, in dynamic scaling of behavioral responses, roles that have conflicting implications for understanding STN function in the context of deep brain stimulation (DBS) therapy. To examine the nature of event-related STN activity and subthalamic-cortical dynamics, we performed primary motor and somatosensory electrocorticography while subjects (n=10) performed a grip force task during DBS implantation surgery. The results provide the first evidence from humans that STN gamma activity can predict activity in the cortex both prior to and during movement, consistent with the idea that the STN participates in both motor planning and execution. We observed that STN activity appeared to facilitate movement: while both movement onset and termination both coincided with STN-cortical phase-locking, narrow-band gamma power was positively correlated with grip force, and event-related causality measures demonstrated that STN gamma activity predicted cortical gamma activity during movement. STN participation in somatosensory integration also was demonstrated by casual analysis. Information flow from the STN to somatosensory cortex was observed for both beta and gamma range frequencies, specific to particular movement periods and kinematics. Interactions in beta activity between the STN and somatosensory cortex, rather than motor cortex, predicted PD symptom severity. Thus, the STN contributes to multiple aspects of sensorimotor behavior dynamically across time.

## Introduction

Understanding the network-level encoding of movement is critical for improving basal ganglia-thalamocortical circuit models, for developing closed-loop deep brain stimulation paradigms, and for developing brain-computer interfaces that combine cortical and subcortical signals to control neuroprosthetic devices. Movement-related information transfer in the cortex is thought to be reflected in the coupling of local field potentials (LFP), where event-related broadband gamma activity is thought to index population firing of local principal neurons (Manning et al., 2009). Slower oscillations, such as those in the beta frequency band (13–30Hz), represent rhythmic fluctuations of neuronal excitability that may serve to coordinate information transfer between regions (Jensen et al., 2005; Yamawaki et al., 2008), including within the basal ganglia-thalamocortical network. How specific cortical-subcortical interactions encode specific aspects of movement, is not well understood. Invasive recordings in subjects implanted with deep brain stimulation (DBS) electrodes in the basal ganglia represent the optimal paradigm for obtaining this information from humans.

The STN is a primary DBS target used in the treatment of Parkinson’s disease (PD) that receives cortical input directly from the frontal lobe and indirectly through the striatum (Afsharpour, 1985; Haynes and Haber, 2013; Nambu et al., 1997; Parent and Hazrati, 1995). Studies examining LFP recorded from the STN with simultaneous recordings from scalp EEG or MEG demonstrated that gamma and beta frequency band activity is coherent between cortical sources and the STN (Fogelson et al., 2006; Litvak et al., 2012; Williams et al., 2002), a relationship modulated both by levodopa (Williams et al., 2002) and movement (Lalo et al., 2008; Litvak et al., 2012). Cortical activity in the beta band has been suggested to drive activity in the subthalamic nucleus at rest (Lalo et al., 2008; Litvak et al., 2011, 2012; Oswal et al., 2016; Williams et al., 2002), a causal interaction that is attenuated with movement (Litvak et al., 2012). EEG and MEG recordings, however, do not have sufficient spatial resolution for reliably localizing the specific anatomical source of cortical oscillations at the level required for precise causal analyses, for instance differentiating activity originating in primary motor (M1) from that in primary sensory (S1) cortex. These methods also have been unable to identify gamma-range causal interactions during movement (Lalo et al., 2008; Litvak et al., 2012), likely due to low signal to noise ratio.

The recent adaptation of electrocorticography to intraoperative neurophysiology research in subjects undergoing DBS safely allows for invasive recording of simultaneous STN-somatomotor cortex LFP activity during the awake portion of the procedure, while patients perform behavioral tasks (Panov et al., 2016). To define the temporal dynamics of movement-related information transfer between the cortex and STN, we employed intracranial recording techniques (Crowell et al., 2012) to collect STN LFPs, simultaneous with electrocorticography (ECoG) from primary motor cortex (M1) and primary somatosensory cortex (S1), during a hand grip task (Fig. 1A). In addition to standard measures of movement-related spectral dynamics, including power, phase-amplitude coupling and phase coherence, we performed event-related causality (ERC) analysis, which uses a multivariate autoregressive technique to estimate causality in multichannel data (Korzeniewska et al., 2008), to define the temporal evolution of task-related causal influences in both beta and gamma frequency ranges. Finally, we determined which of these interactions were correlated with task-related motor performance and motor symptoms of PD.

**Figure 1.**
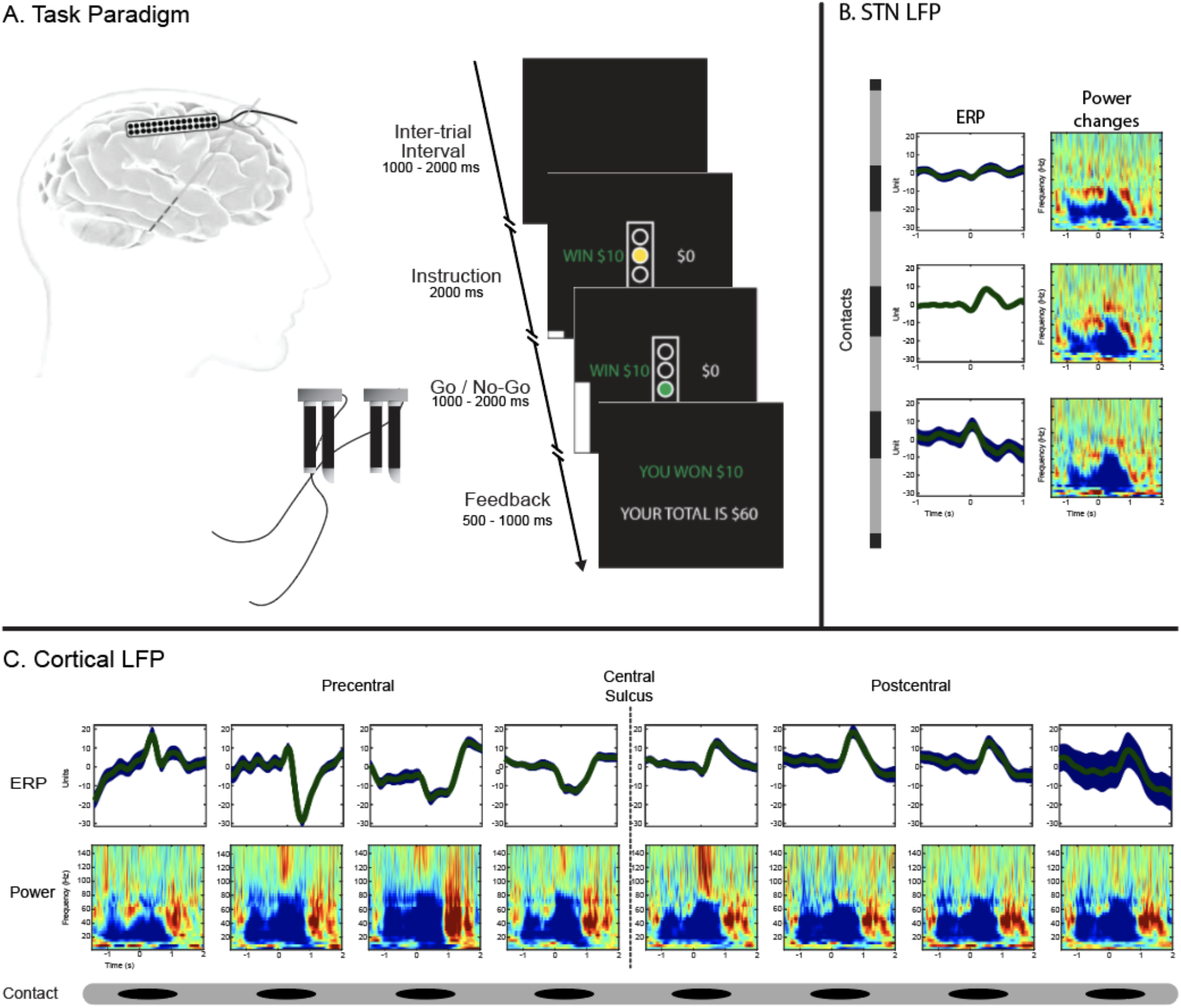
Schematic of experimental design. A simplified schematic of the cue screens, intervals, and recording paradigm is shown in Panel A. Subjects squeezed grip-force transducers in response to visual stimuli. The inter-trial interval duration was randomized on a per trial basis, while the Go/No-Go and feedback durations were adjusted per subject after a training session established how quickly the subject performed the task. Simultaneous recordings using implanted subdural electrodes over motor and sensory cortices (Panel C) and electrodes on the STN DBS lead (Panel B) demonstrate that movement was associated with an ERP in both the STN and cortical areas. The mean ERP (green) and standard deviation (blue) at each recoding contact, are shown from a representative patient. Movement induced a similar pattern of spectral changes at all recordings locations (power modulation shown as z-scores relative to a 1-second baseline at each recording site).

## Methods

### Subjects

Study subjects were recruited from a population of patients with PD scheduled to undergo DBS implantation. Subjects were recommended for surgery by a multidisciplinary review board based on standard clinical indications and inclusion/exclusion criteria. Subject demographics are shown in Table 1. Informed consent was obtained prior to surgery in accordance with a protocol approved by the Institutional Review Board of the University of Pittsburgh (IRB Protocol # PRO13110420).

**Table 1.** Subject demographics

| Subject | Age | Gender | Disease Duration (years) | MMSE | Rest tremor |  | Action tremor |  | Finger Taps |  | Hand Grip |  | RAM |  |
| --- | --- | --- | --- | --- | --- | --- | --- | --- | --- | --- | --- | --- | --- | --- |
|  |  |  |  |  | R | L | R | L | R | L | R | L | R | L |
| P1 | 44 | M | 10+ | 29/30 | 3/3 | 3/3 | 4/3 | 4/3 | 0/0 | 1/1 | 0/0 | 1/1 | 0/0 | 1/1 |
| P2 | 52 | M | 12 | 27/30 | 1/0 | 3/0 | 1/0 | 1/0 | 2/0 | 3/1 | 2/0 | 3/1 | N/A | N/A |
| P3 | 59 | M | 7 | 29/30 | 2/0 | 3/0 | 1/0 | 1/0 | 2/1 | 3/2 | 2/1 | 2/1 | 2/1 | 2/1 |
| P4 | 54 | F | 28 | 29/30 | N/A | N/A | N/A | N/A | N/A | N/A | N/A | N/A | N/A | N/A |
| P5 | 60 | M | 12 | 30/30 | 2/0 | 2/0 | 1/1 | 2/2 | 1/0.5 | 1/0.5 | 1/1 | 1/1 | 1/1 | 1/1 |
| P6 | 46 | M | 4 | 29/30 | 0/0 | 1/0 | 0/0 | 1/0 | 2/1 | 3/1 | 2/1 | 3/2 | N/A | N/A |
| P7 | 53 | M | 10 | 30/30 | 1/0 | 0/0 | 0/0 | 0/0 | 2/1 | 2/1 | 2/1 | 2/2 | 3/1 | 3/2 |
| P8 | 66 | M | 10+ | 30/30 | 3/1 | 3/3 | 2/1 | 3/2 | 3/2 | 3/3 | 3/1 | 3/2.5 | 3/2 | 2/2 |
| P9 | 51 | M | 10+ | N/A | 0/0 | 0/0 | 0/1 | 0/0 | 1/0 | 1/1 | 1/0 | 1/0 | 1/0 | 1/0 |
| P10 | 71 | M | 22 | 30/30 | 2/1 | 0/0 | 0/0 | 0/0 | 3/2 | 2/1 | 3/2 | 3/1 | 3/2 | 3/1 |

### Behavioral Paradigm

The subjects performed a visually cued, instructed delay handgrip task with monetary reward, as previously described (Kondylis et al., 2016). Briefly, a trial began with simultaneous presentation of a yellow traffic light in the center of the screen, and a cue on one side indicating which hand the patient should use for the subsequent response (squeezing the handgrip). The cue remained on the screen for 1000–2000 ms, following which the traffic light changed to either green (Go cue) or red (No-Go cue). A grip force response ≥10% of a previously measured maximum voluntary grip force for ≥100 ms, within 2 seconds of the Go cue onset on the correct side, was considered a successful Go trial. A trial was counted as an error if subjects either did not meet these criteria, or inappropriately squeezed during a No-Go trial. Finally, a feedback message cued the subject to stop squeezing the handgrip and indicated the dollar amount won or lost for the trial, as well as a running balance of winnings. Each trial was followed by a variable inter-trial interval of 500-1000 ms. Subjects performed the task for a cumulative total of 10 to 25 minutes, and only data was only analyzed from subjects who achieved >20 successful trials for each hand. No-Go trials were not analyzed, due to insufficient numbers of No-Go trials for statistical comparison.

### Electrophysiological Recordings

ECoG data was recorded intraoperatively from subjects with movement disorders using a standard 4-contact (n=1), 6-contact (n=5) or 8-contact (n=3) strip electrode (2.3 mm exposed electrode diameter, 10 mm inter electrode distance) (Ad-Tech, Medical Instrument Corporation, Racine, WI), temporarily implanted through the burr hole used for DBS lead implantation as previously described (Crowell et al., 2012; Kondylis et al., 2016). In one subject, a 2×14-contact electrode was used (1.2 mm exposed electrode diameter, 4 mm inter electrode distance). LFPs from the STN were recorded using the clinical DBS lead (model 3389, Medtronic, Minneapolis, MN), except for one subject, in which the STN LFPs were recorded from a ring contact located 3 mm superior to the tip of the 3 microelectrodes used for microelectrode recordings, sampled at 1375 Hz. A referential montage was used with the reference electrode placed in the scalp and a ground electrode placed in the skin overlying the acromion process. Anti-parkinsonian medications were held for at least 12 hours prior to intraoperative testing. LFP data from the lead was obtained after clinical stimulation testing was completed. Subjects were fully awake, without administration of anesthetic agents for at least one hour prior to task performance. No medication was given during task performance. Seven subjects underwent unilateral recordings from the left side during task performance, and three subjects performed the task on both side. ECoG and STN signals were filtered (0.3Hz – 7.5kHZ), amplified and digitized at 30 kHz using a Grapevine neural interface processor (Ripple Inc., Salt Lake City, UT).

The task paradigm was implemented using Psychophysics Toolbox (Brainard, 1997) on a portable computer. Force signals from the handgrips and triggers marking the presentation of visual cues were digitally recorded simultaneously with the ECoG signals. Movement onset was calculated offline by smoothing force signals (15 ms running average) and using a 50 N/s threshold to detect changes in the rate of force generation. The onset of derived grip force data was used to segment ECoG data, and the triggers were used to isolate successful, contralateral Go trials.

### Electrode Localization

Temporarily implanted subdural electrode strips were localized using a method to align pre-operative MRI, intraoperative fluoroscopy and post-operative CT (Randazzo et al., 2016). Briefly, the CT and MRI were coregistered using mutual information in the SPM software package and rendered into 3-D skull and brain surfaces using Osirix and Freesurfer software (Dale et al., 1999), respectively. These surfaces and the fluoroscopy image were then loaded into a custom Matlab user interface and aligned using common landmarks: stereotactic frame pins, implanted depth electrodes, and skull outline. The parallax effect of the fluoroscopic images was accounted for using the measured distance from the radiation source to the subject’s skull. Following surface-to-fluoroscopic image alignment, a 3-D location for each electrode was projected from the fluoroscopic image onto the cortical surface. Based upon the cortical parcellation for each subject’s anatomy (Desikan et al., 2006), each electrode was assigned to a cortical gyrus. Electrodes were then grouped into anatomical Regions of Interest (ROIs).

### Data Preprocessing

All electrophysiological data were preprocessed in MATLAB using custom scripts. First, DC offsets were removed from each channel. Line noise at 60 Hz and its harmonics was removed using a notch filter (MATLAB function *idealfilter*). Next, the data was low-pass filtered at 400 Hz using zero-phase finite impulse response (FIR) filters custom designed in Matlab. The data was resampled to a sampling frequency of 1200 Hz in two steps. Channels with extensive artifact from movement, powerline or environmental sources were visually identified and removed from further analysis. To minimize noise and ensure recordings were comparable across acquisition environments, LFP signals were re-referenced offline to a bipolar montage for the STN channels, and to common average reference for the cortical ECoG channels. All trial epochs were visually inspected for any residual artifact and trials with any contaminated segments were rejected.

### Electrode Selection

Electrode contacts were selected for further analysis based on anatomical and functional considerations. First, only the cortical electrodes localized to M1 (precentral gyrus) and S1 (postcentral gyrus) were included. Event-related potentials (ERPs) centered on movement onset were then used as an independent physiologic measure to select electrode contacts for subsequent time-frequency analyses. Briefly, the mean voltage during baseline was subtracted from the ERP and each time point was tested against zero using a t-test. Surrogate ERPs were constructed from randomly sampled time points and tested the same way. An electrode was considered functionally activated if it showed a cluster of time points with a t-statistic sum greater than the 95% of the cluster sums in the null distribution from the surrogate ERPs (Groppe et al., 2011).

### Spectral Analysis

#### Event-Related Amplitude Modulation

Channel data was temporally convolved with complex Morlet wavelets to obtain the instantaneous spectral components of the signal (Tallon-Baudry et al., 1999). The wavelet transform was calculated in steps of 2 Hz from 10 to 34 Hz for the beta band and in steps of 5 Hz from 50 to 150 Hz for the gamma band. The modulus of the complex signal, representing the analytical amplitude for each band, was divided into 3.5 s epochs surrounding movement onset (1.5 s pre-movement and 2 s of post-movement).

#### Phase-Amplitude Coupling

First, channel data was bandpass filtered using a FIR filter [EEGLAB (Delorme and Makeig, 2004)] into 0.5 Hz bands from 10 to 34 Hz in 0.5 Hz steps for subsequent phase estimation and into 5 Hz bands from 60 to 400 Hz in 5 Hz steps for subsequent amplitude estimation. Second, instantaneous phase and amplitude were estimated using the Hilbert transform. The data was then split into trial epochs as described above and further windowed into 0.1 s non-overlapping windows (35 total windows from −1.5 s to 2 s relative to movement onset). Phase and amplitude was concatenated across trials in each window to create a single continuous phase signal and a single continuous amplitude signal for each electrode across each frequency. The strength of PAC was measured using the Modulation Index (MI) (Tort et al., 2008) calculated with 15 phase bins (24° bin width). We previously demonstrated that the total magnitude of phase-amplitude coupling and the proportion of electrodes recording significant phase-amplitude coupling from high density ECoG strip electrodes fell within the respective ranges for recordings from standard size electrode strips (Kondylis et al., 2016).

#### Inter-regional Phase-locking

Phase-locking value (PLV) measures inter-regional synchrony by quantifying the consistency in phase difference across trials relative to a stimulus (Lachaux et al., 1999). While both amplitude and phase covariance contribute to classical measures of coherence, PLV relies only on the phase information and is thus agnostic to power covariation. To compute PLV, channel data was temporally convolved with complex Morlet wavelets to obtain the instantaneous spectral components of the signal (Tallon-Baudry et al., 1999). The wavelet transform was calculated between 10 and 34 Hz in steps of 1 Hz for the beta band and between 60 and 115 Hz in steps of 2 Hz for the gamma band. The real and imaginary component of each time-frequency point was divided by the modulus of the vector to generate a signed, unit-length, complex-valued time series. Trial epochs were constructed as above. The PLV between two channels was calculated by taking the modulus of the mean of the complex-valued product of multiplying trial epochs from one channel with the complex conjugate of a second channel. The PLV for frequency band f between sources i and j was defined as:

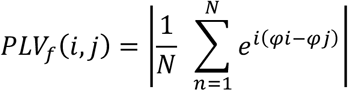

where N is the number of trials and ϕ represents the phase of the data at the wavelet frequency f for sources i, resulting in a frequency-specific PLV, having a value from [0, 1], where 0 represents total phase independence and 1 means that all phase values are equal between the two signals.

### Event-Related Causality

ERC is based on the concept of Granger causality (Granger, 1969) where for signal Y to be considered causally influenced by signal X, knowledge of X’s past has to significantly improve the prediction of Y. ERC uses a multivariate autoregressive (MVAR) model that enables estimation of causality in multichannel data in short windows (Ding et al., 2000). ERC estimates only direct causal influences by using short-time direct Directed Transfer Function (SdDTF) (Korzeniewska et al., 2008) and employs a semiparametric regression model to investigate statistically significant event-related changes in effective connectivity across time (Korzeniewska et al., 2008, 2011). This family of methods has a specific advantage over other effective connectivity measures, in that it mitigates the effects of volume conduction by minimizing the effect of zero-phase-delay conduction, and it has been validated in previous ECoG work (Blinowska et al., 2010; Flinker et al., 2015; Korzeniewska et al., 2011; Nishida et al., 2017).

Briefly, for each subject the signal was bandpassed between 65-115 Hz an FIR filter as implemented in EEGLAB (Delorme and Makeig, 2004). The signals were then downsampled to 400 Hz and segmented into 2048 sample (5.12 s) epochs centered on the cue stimulus, command stimulus and the response time. SdDTF was calculated in short windows of 0.3 s, shifted in time by 0.012 s, for multiple realizations of the same stochastic process (many trials/repetitions of the task). Only channels meeting the channel selection criteria described above were included in the model. Multiple trials or task repetitions from the same subject may be treated as repeated realizations of the same stochastic process which is stationary over short periods. ERC values locked to response onset were statistically tested with a baseline distribution from a 1 second precue baseline and Bonferroni-corrected for multiple comparisons. ERC values passing significance were retained in the frequency range of 65–115 Hz for the gamma band and 12–30 Hz for the beta band, normalized within subject and averaged across subjects and anatomical regions. A similar procedure was applied for ERC analysis in the beta frequency band. However, the signal was band-pass filtered between 13–30 Hz and downsampled to 120 Hz. The window used for SdDTF calculation was 0.4 s long and shifted in time by 0.016 s. While ERC values for individual subjects represent statistically significant predictions, we tested whether the group showed consistent predictions at similar time periods. A right-sided t-test was used to test if the group ERC values at each time point were significantly above a mean of zero which represents a null hypothesis that there is no consistent ERC prediction at this time point. A false discovery rate of 5% was used to correct for multiple comparisons. Next, we sought to define time periods when ERC predictions in a certain direction were significantly larger. For each of the nine flow directions, a pairwise comparison was performed between every two flows with a mutual node by performing a t-test at every time point whether the mean ERC values for a flow A was significantly different than the ERC values of flow B. A positive t-statistic means that flow A was larger and a negative value means that flow B was larger. A cluster-based permutation procedure was performed to find the significant time periods by randomly assigning individual ERC value to each flow direction and performing the t-test as above a 1000 times to construct a null distribution based on the maximal cluster t-statistic. A p-value was assigned to the clusters found in the real data based on their location relative to the null distribution.

### Correlation with task-related motor performance

Trial to trial variability in spectral power or phase-locking can reveal task-related motor performance encoding at a cortical site (Cohen and Cavanagh, 2011). Therefore, we correlated spectral power at every time-frequency point with performance across each trial. Prior to averaging across an ROI for a subject, the correlation values were z-score normalized using a surrogate distribution built by shuffling the trial to performance value mapping 1000 times. We tested the group z-scores against zero using a one-way t-test. To define the significant correlations, we adapted previously described methods for control of multiple comparisons (Smith and Nichols, 2009). Briefly, we generated a surrogate distribution by randomly flipping the sign of the time-frequency image for a random number of subjects, effectively substituting that subject with the mean of the surrogate distribution for that subject. We then performed the t-test and repeated this procedure 1000 times. This maintained the autocorrelation inherent in the data. Clusters surpassing a p-value of 0.05 were considered significant. A similar procedure was performed for PLV but instead we assessed circular-linear correlation using (Berens, 2009)(circ_corrcl.m in the CircStat toolbox) which linearizes the phase difference between two channels into its sine and cosine components and calculates a single correlation coefficient between the phase difference values across trials to all performance values at every time-frequency point. Normalization and statistical significance testing was performed as above.

For PAC, the average performance variables for each subject were correlated with the MI values at each phase-amplitude frequency pair using spearman’s correlation in each temporal window. To define significant group correlations, the subject mapping of the average performance variables was shuffled 1000 times and correlated to the MI values. The real and surrogate correlations were z-scored to the mean and standard deviation of the surrogate distribution. Each z-score image was thresholded at a z-value corresponding to a p-value of 0.05. A p-value was assigned to the clusters of correlation in the real data based on its percentile in the distribution of largest cluster z-score sum in the surrogates. The cluster p-values were finally corrected for the number of time windows with a false discovery rate of 5%.

To define epochs of functionally significant correlation in the ERC, the pairing between each subject’s task-related motor performance variable and ERC time course was randomly shuffled 1000 times. The real correlation and surrogates were z-scored to the mean and standard deviation of the surrogate distribution. The timecourse was thresholded at a z-value corresponding to a p-value of 0.05. The sum of z-values of continuous suprathreshold points were compared to the similar points in the surrogate distribution. A p-value was assigned to the epochs of correlation in the real data based on its location relative to this distribution.

Finally, we investigated the effect of disease severity on the measured spectral changes. For each subject, the average z-scored spectral measure at each location was correlated with their respective Unified Parkinson’s Disease Rating Scale (UPDRS) Part Three total score in the OFF-medication state at every time-frequency point (frequency-frequency point for PAC). The data from the two subjects with bilateral recordings was averaged prior to the correlation and one patient was excluded due to the lack of a UPDRS score. To define significant correlations, the subject mapping of UPDRS scores was shuffled 1000 times to construct a surrogate distribution of correlation values. The real and surrogate correlations were z-scored to the mean and standard deviation of the surrogate distribution. Each z-score image was thresholded at a z-value corresponding to a p-value of 0.05. A p-value was assigned to the clusters of correlation in the real data based on its percentile in the distribution of largest cluster z-score sum in the surrogates. We retained clusters with a corresponding p-value less than 0.05.

### Statistical Analysis

Significance at the group level was assessed using paired cluster-based permutation testing (Maris and Oostenveld, 2007). For spectral amplitude and PLV, we used a standard resampling technique to construct a surrogate baseline (Voytek et al., 2010). For spectral amplitude, surrogate trials were constructed by using randomly generated movement onset times to epoch trials as above. Surrogate PLVs were created by shuffling the trials in one channel, thus eliminating the trial dependence between the two channels (Lachaux et al., 1999). This procedure was repeated 1000 times and mean of surrogates was used as the baseline epoch for that channel. The PAC during a 1 s pre-cue baseline was calculated in a similar fashion to trial PAC and used for the permutation testing at the group level. Measures from all the channels within an ROI were averaged on a per subject basis prior to group level statistics.

Briefly, a paired t-test was performed at every time-frequency point (phase-amplitude pair for PAC) for both trial amplitude or phase-locking between the real group data and the group surrogate baseline for a given subject (actual baseline for PAC). The resulting t-statistics map was thresholded at a pixel p-value of 0.05. The sum of the t-statistics within the resulting clusters was calculated. The trial and baseline data were shuffled while retaining the subject mapping between trial and baseline and tested again as described for the real data. This procedure was repeated 10000 times to construct a null distribution based on the maximal cluster t-statistic found at each permutation. A p-value was assigned to the clusters found in the real data based on their location relative to the null distribution. To account for multiple comparisons concerning the number of ROIs and frequency bands, we applied a Holm-Bonferonni correction (Holm, 1979) and only presented data for significant clusters after correction. For the windowed PAC clusters, only clusters with a p-value < 0.05 after a false discovery rate of 5% correction for multiple comparisons were retained. Event timings for significant changes were based on the temporal bounds of the significant clusters.

## Results

### Motor Performance

Ten subjects with PD were studied, 3 of whom performed the task during recording in both hemispheres, resulting in 13 total hemispheric recordings. The subjects’ motor performance fell within the previously published performance range of a partially overlapping (n=3) cohort of subjects performing the same task, whose measures of motor performance we previously reported to not differ from that of subjects with either essential tremor or no movement disorder, in terms of reaction time (t-stat=1.8, p-value=0.08), time-to-peak (t-stat=1.4, p-value=0.18) and peak force (t-stat=0.03, p-value=0.97) (Kondylis et al., 2016). The average movement duration, defined as the period from force inflection to return to baseline, was 1340ms ± 238ms (mean ± std). The evolution of force generation was stereotyped within subjects across trials, reaching peak force at an average of 630 ± 210ms (mean ± std) in relation to movement onset, and returning to baseline on average at 1340ms ± 238ms (mean ± std).

Event-related potentials (ERPs) coincident with the time of movement onset were calculated for all electrode recording channels and this independent measure of neural activation was used to select channels of interest for further analyses (Fig. 1). Twelve out of the thirteen ECoG recordings (9/10 subjects) showed a significant ERP response in both M1 and S1 gyri (1–3 contacts per region) with reversal of the ERP polarity indicating electrode locations on either side of the central sulcus in each case. In the remaining subject, no contacts were localized to S1, but a significant ERP response was found in contacts overlying M1. In the STN, at least one bipolar pair with a significant ERP response was found in 12 of 13 recordings, and the bipolar contact with the largest ERP for each subject was selected for further analysis. The one recording that lacked an ERP response in the STN was excluded from further analysis.

### Event-Related Modulation of Oscillatory Activity

We found that movement was preceded by beta desynchronization across the motor circuit involving M1, S1 and the STN that was followed by an increase in gamma activity centered around movement (Brittain et al., 2014; Brown et al., 2001; Crone et al., 1998a, 1998b). Group-level analysis of spectral power changes during motor planning and execution, demonstrated a common pattern of beta power modulation in each region. An early decrease in power began prior to movement onset and persisted throughout the movement epoch. This movement-related beta desynchronization appeared in close temporal fashion across locations, at −953ms (M1) (cluster bounds: −953 to 805 ms/10–35Hz, cluster p-value 5e^−5^), −937ms (STN) (cluster bounds: −937 to 952 ms/10–35 Hz, cluster p-value 5e^−5^) and −826ms (S1) (cluster bounds: −826 to 979 ms/10–25Hz, cluster p-value 5e^−5^). In all 3 regions, beta band power rebounded around the time of movement termination, at +1157ms (STN) (cluster bounds: +1157 to 2000 ms/12–35 Hz, cluster p-value 0.0031), +1384ms (S1) (cluster bounds: +1384 to 2000ms/10–35Hz, cluster p-value 0.015), and +1479ms (M1) (cluster bounds: +1479 to 2000 ms, cluster p-value 0.031). Gamma modulation also occurred in each region, including S1. A circuit-wide increase in amplitude occurred in the immediate peri-movement period at −163ms (M1) (cluster bounds: −163 to 552 ms/ 65–150Hz, cluster p-value 0.0001), −105ms (S1) (cluster bounds: −105 to 721 ms/60–150Hz, cluster p-value 0.0012), and +250ms (STN) (cluster bounds: +250 to 459 ms/60–100Hz, cluster p-value 0.004). Changes in gamma activity spanned a broadband range in both cortical areas, while that in the STN occurred predominantly in a narrow band centered at 75 Hz (based on the weighted centroid of the cluster), as previously described for the basal ganglia (Jenkinson et al., 2013), with less modulation in broadband gamma (Fig. 2).

**Figure 2.**
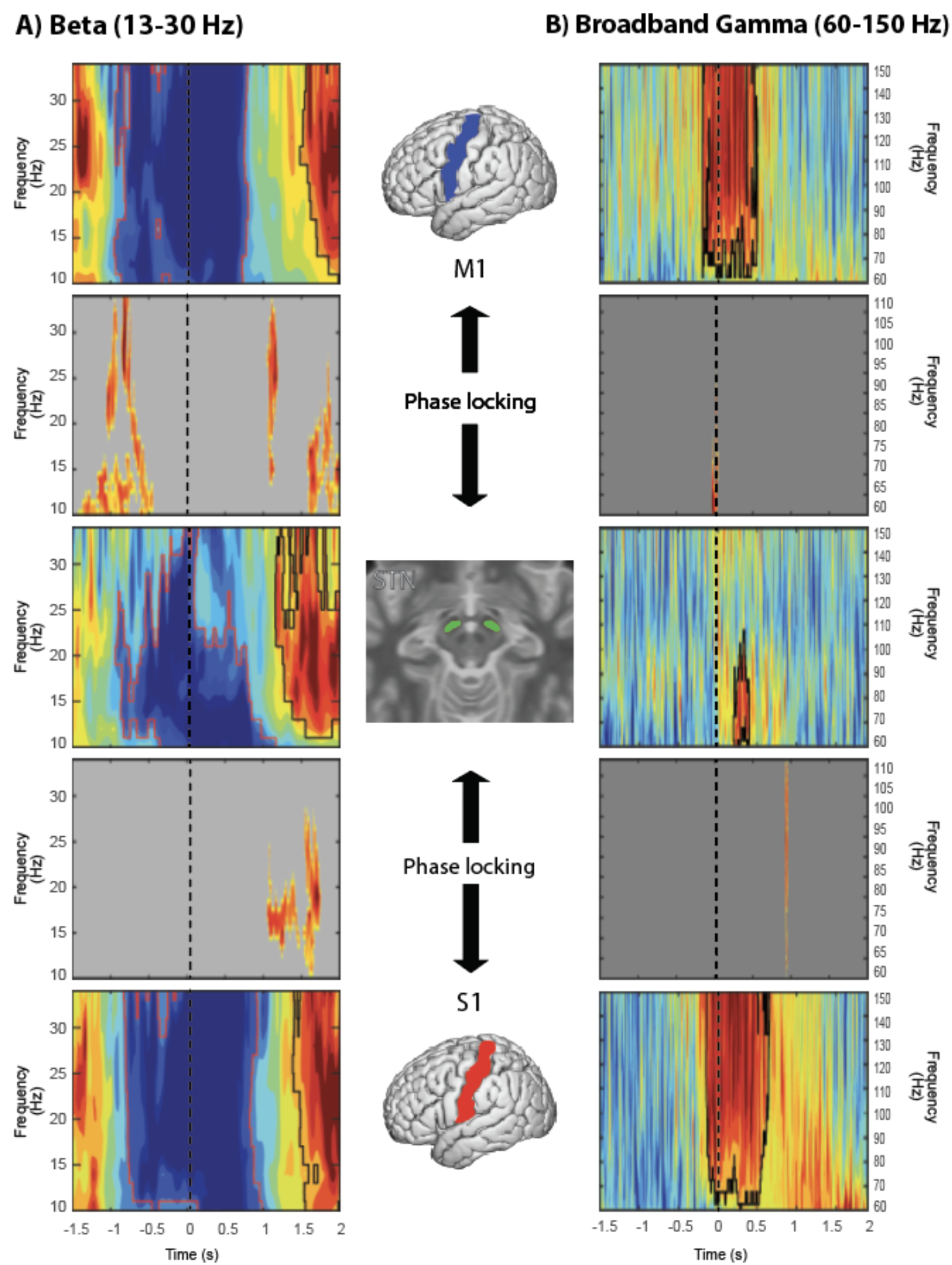
Regional power modulation and interregional functional coupling across the network. Acluster-based permutation test was used to define clusters of significant power changes (panels 1,3 and 5) across subjects marked by a black border for significant increases and a red border for significant decreases. Significant functional coupling as measured by interregional phase-locking was define in a similar fashion (panels 2 and 4) with only significant clusters being plotted. Dotted line represents movement onset.

We next investigated phase-amplitude coupling (PAC) in each brain region. Both cortical regions (M1 and S1) exhibited baseline coupling of the phase of beta oscillations to the gamma amplitude from 60–200Hz, while the STN exhibited significant PAC only between 250-400Hz. Movement-related phase-amplitude decoupling was also seen, slightly preceding movement-related beta desynchronization across the network. A significant decrease in PAC prior to movement was observed in each region, first in S1 (−1100ms), followed by the STN (−1000ms) and M1 (−900ms). Similar to beta power changes near movement termination, a rebound phenomenon for PAC was observed across the entire network, first observed in the STN (+1200ms), followed by M1 (+1300ms), and S1 (+1800ms) (Supp Fig.2). Epochs of PAC that primarily involved phase coupling to low-beta frequencies were observed near movement termination, prior to rebound of baseline PAC, in both M1 and S1, a relationship that was not evident for the STN, which exhibited clusters of re-emerging PAC during movement itself. Phase-amplitude decoupling was observed prior to beta desynchronization, precluding the possibility that phase estimation accuracy was affected by low signal-to-noise ratio from loss of beta power, similar to previously reported findings (Kondylis et al., 2016; Miller et al., 2012). Thus, M1, S1 and STN exhibited movement-related decreases in PAC, though the specific frequencies within the gamma band to which beta phase was coupled were different between these locations.

### Inter-regional Phase-locking

We next examined whether concurrent movement-related amplitude and PAC changes were associated with interregional phase-locking between these three brain regions. With respect to beta activity, prior to movement, M1 exhibited epochs of significant beta phase-locking to both S1 (cluster bounds: −1500 to −1095 ms/10−35Hz, cluster p-value:<0.001) (Supp Fig2.A) and the STN (−1400ms) (Fig. 2). Moreover, M1 beta phase-locking to the STN showed two patterns of synchronization: phase locking to low-beta (centered at 13Hz) and a separate cluster of phase locking to high-beta (centered at 26Hz). The low-beta cluster was sustained from (cluster bounds: −1400ms to −450ms/ 10–16Hz, cluster p-value: 0.00075) while the high-beta cluster persisted for a shorter period of synchronization, from −1054ms to −740ms (cluster bounds: −1054 to −1191ms, −855 to − 740ms/19–33Hz, 16–33Hz, cluster p-values: 0.02, 0.00055 respectively). A rebound phenomenon was observed across the entire network (M1 to S1: 870 to 2000 ms/10–35Hz, cluster p-value:<0.001; STN to S1: +1052 to 1747 ms/11–28Hz, cluster p-value:<0.01), with the M1 to STN beta phase-locking again exhibiting both a high beta cluster (1053 to 1191 ms/14-33Hz, cluster p-value:0.0036) and a low-beta cluster (1576 to 2000 ms/10–27Hz, cluster p-value:0.0004). Thus, STN-M1 phase locking at movement termination recapitulated the pattern seen prior to desynchronization.

Examining phase-locking in the gamma band, we found that phase-locking of STN to M1 and S1 bracketed movement onset and termination (Fig. 2). In the peri-movement period, brief phase-locking in the gamma band centered at 68Hz was observed between M1 and the STN (cluster bounds: −45 to 38 ms/60–92Hz, cluster p-value:0.002) that was followed by M1-S1 phase-locking centered at 88Hz (cluster bounds: 34 to 390 ms/60–115Hz, cluster p-value <0.01 corrected). Near movement termination, brief gamma phase-locking again was observed, between S1 and the STN centered at 92Hz (cluster bounds: 926 to 960 ms/62–115Hz, cluster p-value 0.0037) and M1 to S1 centered at 89Hz (cluster bounds: 1025 to 1496 ms/60–115Hz, cluster p-value <0.01 corrected). Although these gamma phase-locking results theoretically could result from an artifact resulting from increased signal to noise ratio (SNR) in the gamma band, the epoch of movement-related of gamma phase-locking, *preceded* the epoch in which a significant gamma amplitude increase occurred. Similarly, the phase-locking between S1 and M1 was similar in both epochs despite a difference in SNR. Thus, STN gamma phase is coordinated with that of M1 in relation to movement onset and is correlated with that in S1 in relation to movement termination.

### Event-Related Causality (ERC)

We next determined the directionality of neural activity propagation between the STN and both M1 and S1 during different movement periods, using ERC. The ERC method estimates the causal influences (in a statistical sense) between networked brain regions. We used ERC to estimate the direction, intensity and temporal course of propagation for either beta or gamma activity between M1, S1 and the STN, compared to causal interactions during baseline. Statistically significant causal interactions were found across the temporal course of movement planning and execution, as might be expected when assaying connectivity across nodes of a loop circuit. To better describe these interactions, we defined four time periods based on the average times our subjects moved: a motor planning period prior to movement, an early movement period from initiation (time zero) until reaching peak force, a late movement period from peak force until movement termination, and a termination period.

Both gamma and beta band activity showed significant causal interactions between all tested locations. M1 and S1 showed significant bidirectional causal interactions in the gamma band throughout the entire trial epoch. Gamma propagation from M1 to S1 peaked at +264ms while gamma propagation from S1 to M1 peaked at +672ms, coinciding with peak movement. No significant difference in ERC magnitude was detected between the two causal directions. Although M1 and STN also showed significant bidirectional causal interactions, they were limited to the motor planning period and early movement and briefly during movement termination. On the other hand, S1 and STN gamma band interactions were more heterogeneous (Supp Fig.5). Comparing significant pairwise interactions, we observed several important patterns. During movement preparation, the strongest propagation in gamma activity was observed from STN to S1 (STN to S1 vs STN to M1: −1008 to −684ms, cluster p-value: 0.048) (Fig. 3A) (STN to S1 vs S1 to STN: −936 to −540ms, cluster p-value: 0.026) (Fig.3B), and subsequently from S1 to M1 (−804 to −468 ms) (S1 to M1 vs S1 to STN: −804 to −468ms, cluster p-value: 0.029) (Fig. 3C). No causal gamma interactions dominated during the movement epochs. Movement termination was associated with gamma propagation from M1 to STN (M1 to STN vs M1 to S1: 1668 to 2000ms, cluster p-value: 0.016) (Fig. 3D).

**Figure 3.**
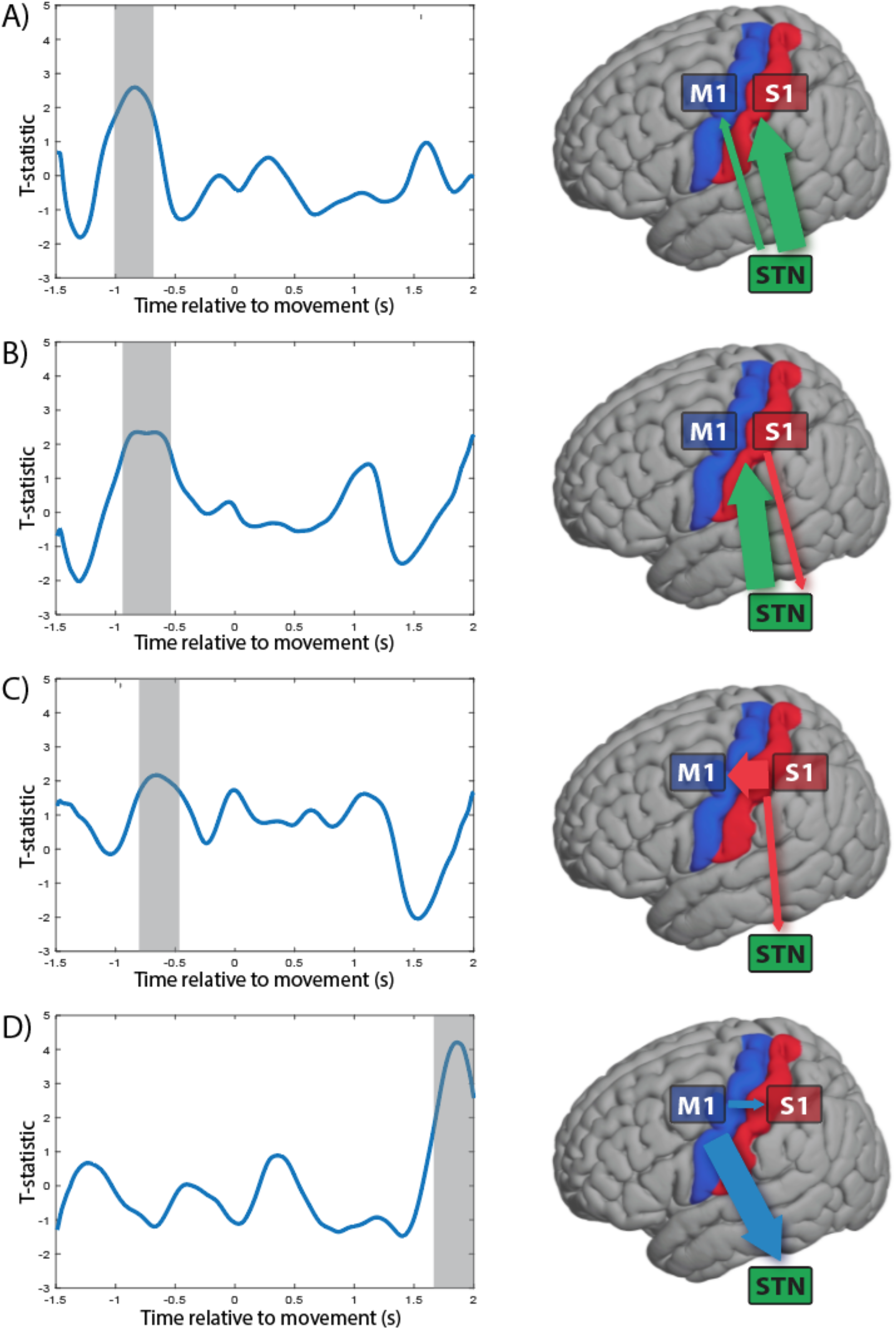
Event-related causal interactions in the gamma band (65–115 Hz). Pairwise comparisons were performed between related causal interactions in the gamma band using a t-test at every time point with the t-statistic plotted as a time course. Significant periods were defined using cluster-based permutation testing; clusters with a p-value less 0.05 compared to a surrogate distribution are marked with gray. During movement planning, three interaction pairs exhibited significant difference STN → S1 vs STN → M1 (A), STN → S1 vs S1 → STN (B), and S1 → M1 vs S1 → STN (C). Movement termination exhibited a significant interaction difference between M1 → STN vs M1 → S1 (D). The causal interactions tested are represented with the arrows on the brain schematic. The color of the arrow corresponds to the area of origin of the causal interaction, and the size depicts the strength of the interaction.

In the beta frequency range, the strongest causal interactions were observed between M1 and the STN, in contrasting patterns. M1 beta activity predicted that in S1 and the STN during movement preparation and again during late movement and termination, but exhibited minimal interactions during movement execution. Beta activity in the STN was predictive of that in M1 and S1 during periods of preparation and early movement, with no outflow interactions near movement termination. S1 beta activity only minimally predicted that in M1 and STN, mainly around movement termination (Supp Fig.4). When comparing the pairwise interactions, we observed the following patterns: during movement preparation and early movement, beta activity in the STN most strongly predicted that in S1 than the reciprocal direction (STN to S1 vs S1 to STN: −828 to −300ms and −124 to 596ms, cluster p-value: 0.0025, <0.001) (Fig. 4A), and even more than M1 (STN to S1 vs STN to M1: −844 to −284ms and −76 to 324ms, cluster p-value: 0.012, 0.03) (Fig. 4B). During movement, S1 received more predictive beta information from the STN than from M1(STN to S1 vs M1 to S1: −252 to 404ms, cluster p-value: 0.038), while at termination, beta information from M1 was most predictive of that in S1 (STN to S1 vs S1 to STN: 1364 to 1700ms, cluster p-value: 0.004) (Fig. 4C).

**Figure 4.**
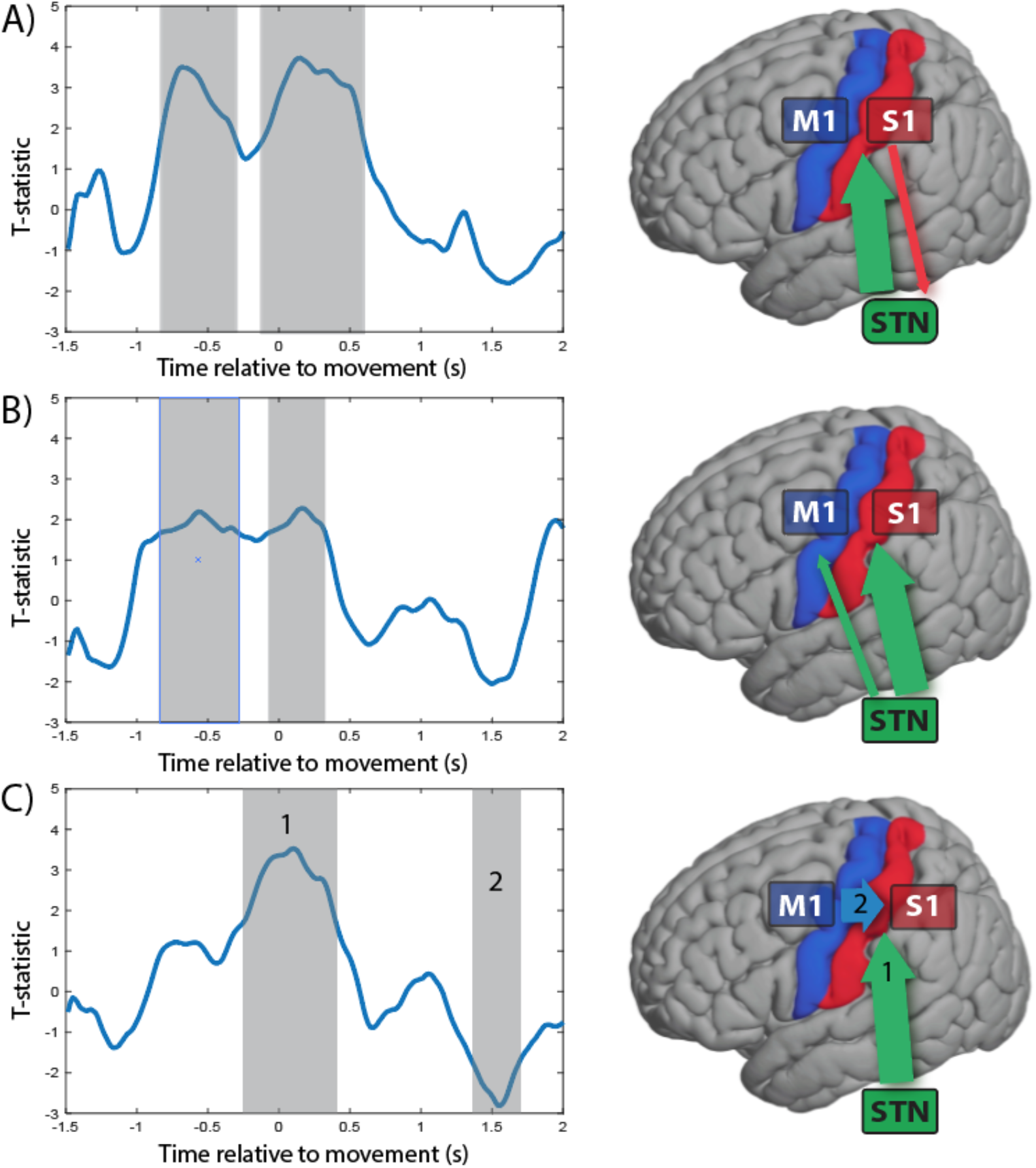
Event-related causal interactions in the beta band (13–30 Hz). Pairwise comparisons were performed between related causal interactions in the beta band using a t-test at every time point with the t-statistic plotted as a time course. Significant periods were defined using cluster-based permutation testing; clusters with a p-value less 0.05 compared to a surrogate distribution are marked with gray. During movement planning and movement execution, two interaction pairs exhibited significant differences, both involving the STN: STN→ S1 vs S1→STN (A), STN→ S1 vs STN→M1 (B). One interaction pair STN → S1 vs M1→S1 (C) exhibited contrasting behavior with larger STN→ S1 flow during movement execution (C, 1) followed by larger M1→S1 during movement termination (C, 2). The causal interactions tested are represented with the arrows on the brain schematic. The color of the arrow corresponds to the area of origin of the causal interaction, and the size depicts the strength of the interaction.

### Encoding of Task-Related Motor Performance

We next determined the extent to which specific features of force generation were encoded in the observed spectral changes. We analyzed four motor performance variables: 1) reaction time, the time between the command stimulus and the force inflection point; 2) peak force, the maximum force generated in a trial, normalized to the session maximum force; 3) peak yank, the maximum of the first differential of the force trace; and 4) peak tug, the maximum of the second differential of the force trace. We found correlations with power and phase based on per patient trial-to-trial variations that are averaged across the cohort, and correlations with PAC and ERC based on cross-subject correlation (note that these measure lack per-trial values). The results are detailed in Table 2.

**Table 2.**
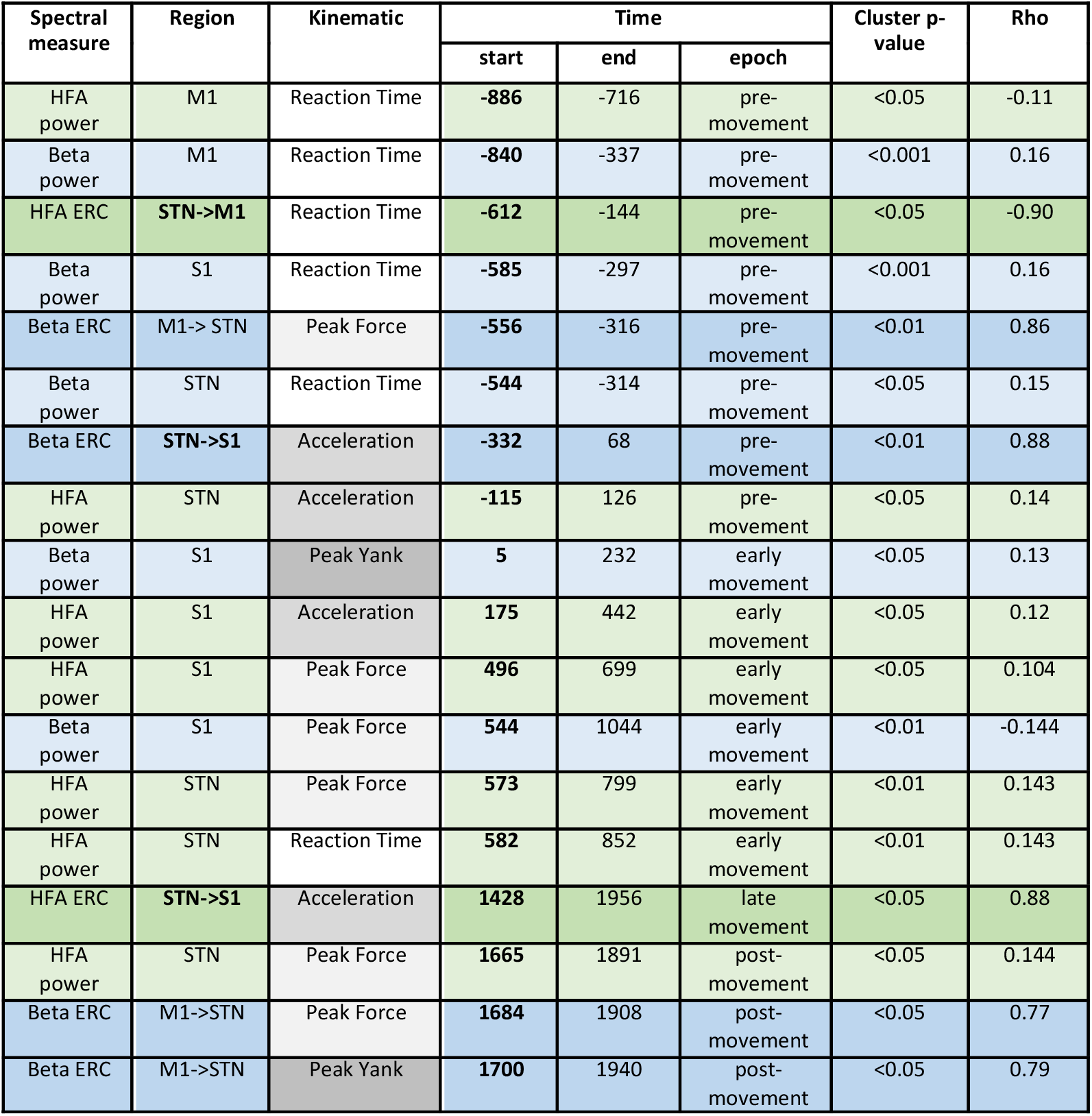
Spectral measures showing correlation to task-related performance

#### Reaction time

In the motor planning period, a greater magnitude of beta power in each region, M1, S1 and the STN, correlated with slower reaction time (Fig. 5A). As expected, gamma power exhibited a contrasting relationship, with greater M1 gamma power in the pre-movement epoch correlated with faster reaction time. Notably, the magnitude of the influence of STN gamma on M1 gamma, prior to movement, was correlated with faster reaction time. A greater magnitude of STN gamma, itself, however, was correlated to slower reaction time, during the early movement period.

**Figure 5.**
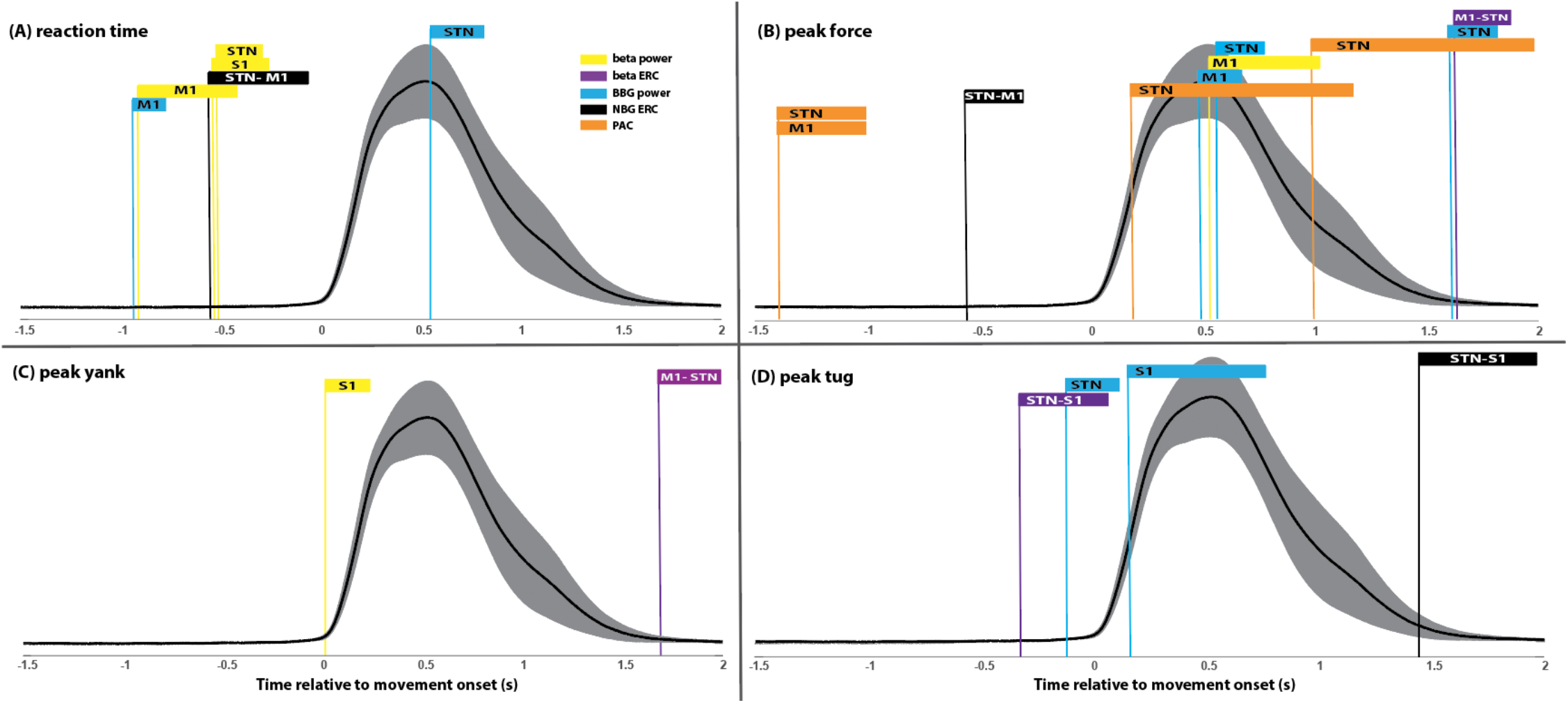
Temporal encoding of force transduction across the network. Group-level, spectral feature-specific epochs exhibiting significant correlation between reaction time (A), peak force (B), peak yank (C) and peak tug (D) are indicated using color bars, preceded by vertical lines intersecting the start time of the statistically significant correlation, on the mean force trace across all subjects (shaded area = 95% confidence interval, BBG = broad band gamma, NBG = narrow band gamma).

#### Peak Force and Yank

In the motor planning period, PAC in M1 and STN were correlated inversely with peak force (Fig. 5B). The strength of M1 beta influence on STN beta activity during this period was correlated with greater peak force. During early movement, M1 beta power itself was correlated with less peak force, while M1 gamma power was correlated with greater peak force. Similarly, the magnitude of STN gamma was correlated with greater peak force, and STN PAC was correlated with less peak force. In the termination epoch, STN gamma again was correlated with the prior generation of higher peak force. M1 influence on STN beta activity in this period was positively correlated with the prior generation of both peak force and peak yank (Fig. 5B).

#### Peak Tug

Interesting dynamics between activity in the STN and S1 were observed in relation to the tug (rate of change in the force per time (yank) measure) (Fig. 5D). STN gamma in the motor planning period was correlated with greater peak tug. STN influence on S1 beta activity during this period also was correlated with greater peak tug. During early movement, S1 gamma power correlated with greater peak tug. Near movement termination, the strength of STN influence on S1 gamma was correlated to the prior presence of greater peak tug.

### Correlation with Parkinson’s Severity

Finally, we assessed the relationship of each of the spectral measures to disease severity. ERC measures in the beta band were the only values found to be significantly correlated to the Unified Parkinson’s Disease Rating Scale (UPDRS) Part 3 score in the OFF-medication state (Fig 6). Specifically, the strength of bidirectional interactions between STN and S1 during the motor planning epoch were negatively correlated with the UPDRS score. During early movement, the strength of beta propagation from M1 to S1 was correlated inversely with UPDRS score. Finally, stronger beta propagation from S1 to STN during movement termination was correlated with lower UPDRS scores.

**Figure 6.**
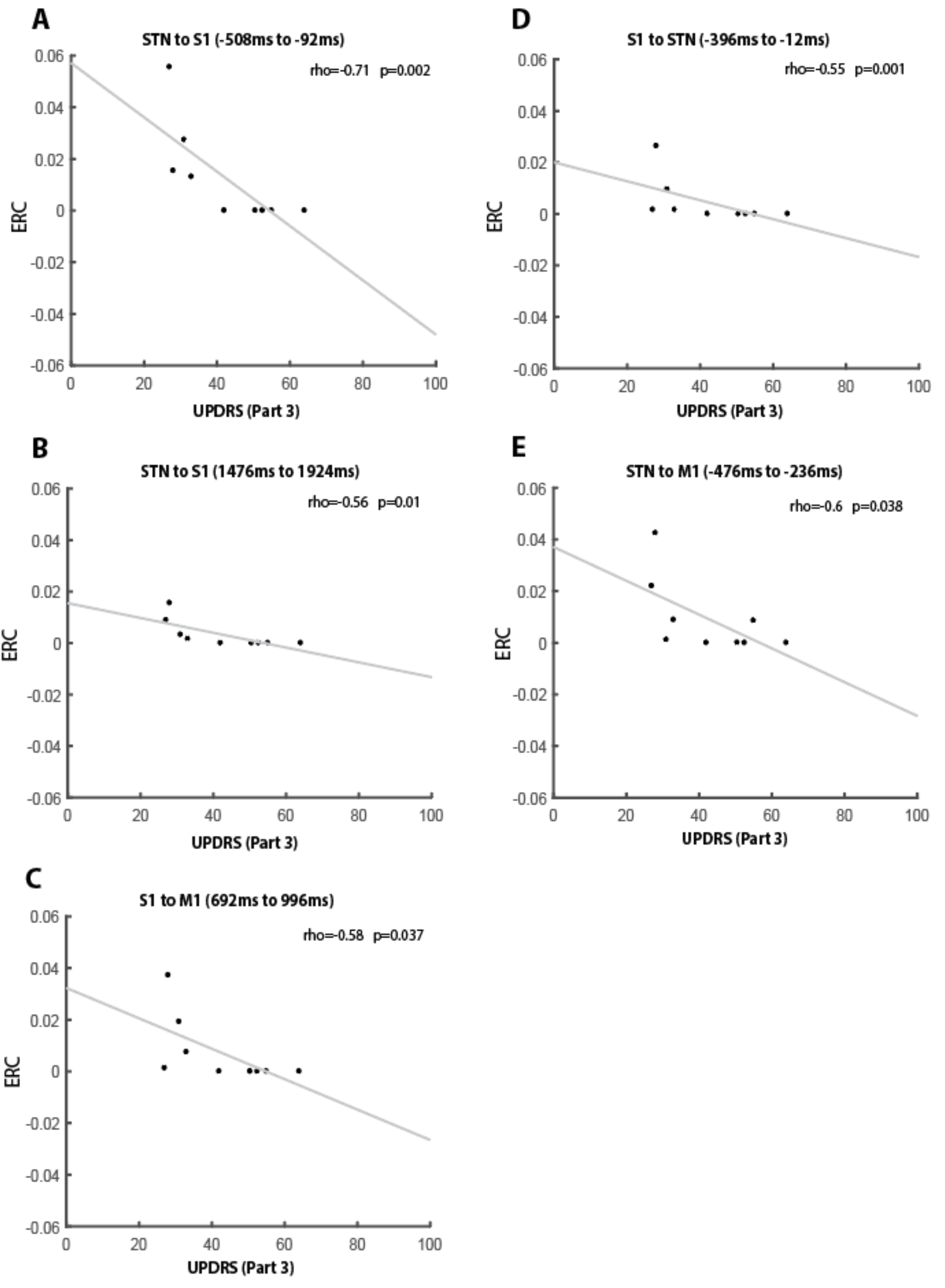
Event-related interactions with significant correlations to UPDRS score. A cluster-permutation test was used to define epochs of significant correlation between an interregional causal interaction and UPDRS scores. Scatter plots show the UPDRS scores for each subject and the mean ERC flow value during significant epochs. The plotted line is the least square line fitted to the data. The average cluster Spearman rho value and p-value of the cluster are represented for each interaction.

## Discussion

We employed intracranial recordings within the cortical-subthalamic network to determine the timing and causal nature of LFP interactions that encode features of force generation. The results provide the first evidence from humans that STN gamma activity can predict activity in the cortex both prior to and during movement, consistent with the idea that the STN participates in both motor planning and execution. Additionally, we demonstrated that S1 plays a significant role in this process, as the STN exhibited distinct interactions with sensory cortex that were specific to certain movement periods and features. Beta interactions between STN and S1, but not M1, predicted disease severity, demonstrating the importance of subcortical integration of sensory information in movement encoding.

### Motor Planning

The STN has been proposed to have a role in motor planning, for instance in preventing or stopping an action, including inhibiting responses during the computation of a decision threshold (Fischer et al., 2017; Herz et al., 2016). For example, gamma power and decreases in STN-cortical synchrony have been correlated with successful stopping action (Fischer et al., 2017), and trial-wise variation in delta-theta oscillatory activity in the STN has been suggested to reflect decision thresholds (Herz et al., 2016). Our use of ERC did reveal that STN activity prior to movement onset contained some information about subsequent force generation. First, the strength of the influence of STN gamma on M1 gamma, in the ~600ms epoch prior to movement, was correlated with faster reaction times. Second, the strength of the influence of STN beta activity on S1 beta activity, in the ~300ms prior to movement, correlated with tug (the rate of change in applied force per unit time). Neither of these results, however, is consistent with the idea that the STN puts a “hold” on cortical action signals (Alegre et al., 2013; Aron and Poldrack, 2006), although they also do not preclude the STN participating in suppressing an already initiated manual response (No-Go trials in this cohort were not analyzed due to insufficient trial number). Our findings better align with the idea that STN gamma activity reflects dynamic processing that supports flexible motor control (Fischer et al., 2017). Although pre-movement beta power and PAC in the STN were correlated with slower reaction times and less peak force generation, we interpret this finding as an indication that excessive beta-range oscillatory activity throughout the network represents a burden of maintenance activity that must be overcome for movement initiation (Miller et al., 2012), since pre-movement M1 and S1 beta power also were each correlated with slower reaction times and the strength of M1 PAC was negatively correlated with peak force.

### Movement Facilitation

Movement-related information transfer may occur through increased gamma synchrony that emerges as the neuronal populations are released from beta entrainment. Recent work supports a model in which cortical beta originates from bursts of synchronous synaptic drivers (Sherman et al., 2016), where the strength of this synchrony is reflected in the waveform properties of the LFP, and increased synchrony lads to non-sinusoidal beta waves, indexed by increased apparent PAC (Cole et al., 2017). The exaggerated PAC seen in PD, therefore, may index over synchronization of M1 input, in which cortical processing is hijacked, hampering information flow (Cole et al., 2017). Consistent with this model, we observed that when beta synchrony and PAC decrease prior to movement, gamma phase synchrony and the ability of STN gamma activity to predict M1/S1 gamma activity increases, prior to movement onset. Similarly, we previously reported that the pattern of STN spike to cortical phase-locking often is modulated before movement (Lipski et al., 2017).

The switch from the resting beta-dominant state to the pro-kinetic gamma-dominant state was marked by a switch from bidirectional causal influence between M1 and STN beta activity (Fig. 4A) to a unidirectional influence of STN beta on cortex, just prior to movement onset (Fig. 4B). The loss of the ability of M1 activity to predict STN activity may be an indication that the predominant input to the STN at this point is not M1, but more likely premotor cortical regions, either via hyperdirect cortical afferents or through the striatal indirect pathway. STN-M1 oscillatory phase alignment in the high-beta range preceded both the onset and offset of movement-related desynchronization. Bracketing of the movement period by coordinated activity between the cortex (M1 and S1) and STN also was evident in the gamma frequency range. STN gamma phase-locking that was specific to M1 followed STN-M1 beta phase-locking and preceded movement initiation. STN gamma phase-locking then preceded the rebound of STN-M1 beta phase-locking and movement termination, but was specific to S1, suggesting that interactions between the STN and S1 are important for stopping an ongoing movement.

Furthermore, our data demonstrate that narrow band gamma oscillations are used to coordinate not only primary motor and subcortical activity (Litvak et al., 2012), but also sensory cortical and subcortical interactions. Across the entire cohort gamma power increases preceded movement in M1 and S1, but occurred after movement onset in the STN, consistent with the idea that the basal ganglia modulate ongoing movements (Mazzoni et al., 2007)(Turner and Desmurget, 2010). We showed that movement is supported by coherent narrow band gamma rhythms across each region in the circuit, which were associated with bidirectional causal interactions between each node. Our findings suggest that online adjustment of motor plan execution is supported by both cortical-cortical and cortical-subcortical feedback through these narrow band gamma rhythms, in line with previous data suggesting their role in facilitating cognitive processing in connected systems (Bosman et al., 2012; Buzsáki and Wang, 2012; Litvak et al., 2012; Montgomery and Buzsáki, 2007; Roberts et al., 2013; Siegel et al., 2008; Womelsdorf et al., 2006).

Our approach additionally revealed that gamma power in the STN also encodes force, in the same temporal window as M1 gamma. The magnitude of STN gamma power near the time of peak force, and the strength of the gamma causal interactions from STN to M1, also positively scaled with reaction time, consistent with the pro-kinetic properties of this narrow band gamma oscillation in the motor circuit (Jenkinson et al., 2013). Previous reports exploring evidence for human STN contributions to movement gain that showed only trends (Tan et al., 2013) or the requirement of auditory cues (Anzak et al., 2012) relied on calculating the average power change rather than the trial-to-trial variability reflected in our analyses.

### Somatosensory Integration

Traditional models of basal ganglia-cortical loop function do not address the role of primary sensory cortex in cortical-subcortical communication. The gating and conversion of sensory information into a form relevant for guiding movement, however, is one mechanism through which the basal ganglia may participate in motor control (Lidsky et al., 1985). Somatosensory cortex projects to the striatum in nonhuman primates (Flaherty and Graybiel, 1991, 1993, 1995), although only in rodents has the STN been reported to receive direct primary somatosensory inputs (Canteras et al., 1990). The high spatiotemporal resolution of ECoG allowed us to demonstrate the independent contributions of interactions with sensory cortex to the encoding of movement features. During the pre-movement epoch, both beta and gamma activity propagated from the STN to S1. S1 interactions were correlated with greater peak tug, indicating critical time points when sensory cortex participates in the scaling of movement. Positive correlations also were observed between the magnitude of S1 gamma power and both yank and tug in the early-movement period. ERC revealed that S1 likely influences STN activity indirectly, following the integration of sensory information in M1, consistent with studies of beta band interactions during static limb position that suggest S1 to M1 propagation (Brovelli et al., 2004; Witham et al., 2010) and SMA/M1 to STN causal influence (Lalo et al., 2008; Litvak et al., 2011, 2012; Oswal et al., 2016; Williams et al., 2002). Recent evidence that spike firing in the STN is entrained to S1 LFPs (Lipski et al., 2017) further supports the presence of functional connectivity between the two regions.

Of all the spectral measures analyzed in this data set, only ERC measures in the beta band were significantly correlated to PD motor symptom severity in the OFF-medication state. Motor symptoms were associated with: 1. the magnitude of bidirectional interactions between STN and S1 during the motor planning period, 2. the magnitude of beta activity propagation from M1 to S1 during early movement, and 3. the magnitude of beta activity propagation from S1 to STN during movement termination. These results support the concept that abnormal sensory integration underlies some PD symptoms (Abbruzzese and Berardelli, 2003).

## Conclusions

This study provides the first evidence from humans that STN gamma activity can predict activity in the cortex both prior to and during movement. In addition, we utilized a novel combination of technical and computational methods to demonstrate that STN activity is associated with the facilitation of force transduction, that the STN participates in somatosensory integration, and that interactions between the STN and somatosensory cortex predict PD symptom severity. These findings have important implications for understanding basal ganglia-cortical circuit disorders and designing novel therapeutic brain stimulation strategies.

## Acknowledgements

Funding support from National Institute of Neurological Disorders and Stroke (U01NS098969) and the Walter Copeland Fund of the Pittsburgh Foundation. The authors declare no competing financial interests.

## References

Abbruzzese, G., and Berardelli, A. (2003). Sensorimotor integration in movement disorders. Mov. Disord. 18, 231–240.

Afsharpour S. (1985). Topographical projections of the cerebral cortex to the subthalamic nucleus. J. Comp. Neurol. 236, 14–28.

Alegre, M., Lopez-azcarate, J., Obeso, I., Wilkinson, L., Rodriguez-oroz, M.C., Valencia, M., Garcia-garcia, D., Guridi, J., Artieda, J., Jahanshahi, M., et al. (2013). The subthalamic nucleus is involved in successful inhibition in the stop-signal task: A local field potential study in Parkinson’s disease. Exp. Neurol. 239, 1–12.

Anzak, A., Tan, H., Pogosyan, A., Foltynie, T., Limousin, P., Zrinzo, L., Hariz, M., Ashkan, K., Bogdanovic, M., Green, A.L., et al. (2012). Subthalamic nucleus activity optimizes maximal effort motor responses in Parkinson’s disease. Brain 135, 2766–2778.

Aron, A., and Poldrack, R. (2006). Cortical and Subcortical Contributions to Stop Signal Response Inhibition: Role of the Subthalamic Nucleus. J Neurosci 26, 2424–2433.

Berens P. (2009). CircStat: A MATLAB toolbox for circular statistics. J. Stat. Softw. 31, 1–21.

Blinowska, K., Kus, R., Kaminski, M., and Janiszewska, J. (2010). Transmission of brain activity during cognitive task. Brain Topogr. 23, 205–213.

Bosman, C.A., Schoffelen, J.-M., Brunet, N., Oostenveld, R., Bastos, A.M., Womelsdorf, T., Rubehn, B., Stieglitz, T., De Weerd, P., and Fries, P. (2012). Attentional stimulus selection through selective synchronization between monkey visual areas. Neuron 75, 875–888.

Brainard D. (1997). The psychophysics toolbox. Spat. Vis.

Brittain, J.-S., Sharott, A., and Brown, P. (2014). The highs and lows of beta activity in cortico-basal ganglia loops. Eur. J. Neurosci. 39, 1951–1959.

Brovelli, A., Ding, M., Ledberg, A., Chen, Y., Nakamura, R., and Bressler, S.L. (2004). Beta oscillations in a large-scale sensorimotor cortical network: directional influences revealed by Granger causality. Proc. Natl. Acad. Sci. U. S. A. 101, 9849–9854.

Brown, P., Oliviero, A., Mazzone, P., Insola, A., Tonali, P., and Lazzaro, V. Di (2001). Dopamine Dependency of Oscillations between Subthalamic Nucleus and Pallidum in Parkinson’s Disease. 21, 1033–1038.

Buzsáki, G., and Wang, X.-J. (2012). Mechanisms of gamma oscillations. Annu. Rev. Neurosci. 35, 203–225.

Canteras, N.S., Shammah-Lagnado, S.J., Silva, B.A., and Ricardo, J.A. (1990). Afferent connections of the subthalamic nucleus: a combined retrograde and anterograde horseradish peroxidase study in the rat. Brain Res. 513, 43–59.

Cohen, M.X., and Cavanagh, J.F. (2011). Single-trial regression elucidates the role of prefrontal theta oscillations in response conflict. Front. Psychol. 2, 30.

Cole, X.S.R., Van Der Meij R., Peterson, E.J., De Hemptinne C., Starr, X.A., and Voytek, X.B. (2017). Nonsinusoidal Beta Oscillations Reflect Cortical Pathophysiology in Parkinson’s Disease. 37, 4830–4840.

Crone, N.E., Miglioretti, D.L., Gordon, B., Sieracki, J.M., Wilson, M.T., Uematsu, S., and Lesser, R.P. (1998a). Functional mapping of human sensorimotor cortex with electrocorticographic spectral analysis. I. Alpha and beta event-related desynchronization. Brain 121 (Pt 1, 2271–2299.

Crone, N.E., Miglioretti, D.L., Gordon, B., and Lesser, R.P. (1998b). Functional mapping of human sensorimotor cortex with electrocorticographic spectral analysis. II. Event-related synchronization in the gamma band. Brain 121 (Pt 1, 2301–2315.

Crowell, A.L., Ryapolova-Webb, E.S., Ostrem, J.L., Galifianakis, N.B., Shimamoto, S., Lim, D. a, and Starr, P. a (2012). Oscillations in sensorimotor cortex in movement disorders: an electrocorticography study. Brain 135, 615–630.

Dale, A.M., Fischl, B., and Sereno, M.I. (1999). Cortical surface-based analysis. I. Segmentation and surface reconstruction. Neuroimage.

Delorme, A., and Makeig, S. (2004). EEGLAB: an open source toolbox for analysis of single-trial EEG dynamics including independent component analysis. J. Neurosci. Methods134, 9–21.

Desikan R.S., Sgonne, F., Fischl, B., Quinn, B.T., Dickerson, B.C., Blacker, D., Buckner, R.L., Dale, A.M., Maguire, R.P., Hyman, B.T., et al. (2006). An automated labeling system for subdividing the human cerebral cortex on MRI scans into gyral based regions of interest. Neuroimage 31, 968–980.

Ding, M., Bressler, S.L., Yang, W., and Liang, H. (2000). Short-window spectral analysis of cortical event-related potentials by adaptive multivariate autoregressive modeling: data preprocessing, model validation, and variability assessment. Biol. Cybern. 83, 35–45.

Fischer, P., Pogosyan, A., Herz, D.M., Cheeran, B., Green, A.L., Fitzgerald, J., Aziz, T.Z., Hyam, J., Little, S., Foltynie, T.,et al. (2017). Subthalamic nucleus gamma activity increases not only during movement but also during movement inhibition. Elife 6.

Flaherty, A.W., and Graybiel, A.M. (1991). Corticostriatal transformations in the primate somatosensory system. Projections from physiologically mapped body-part representations. J. Neurophysiol. 66, 1249–1263.

Flaherty, A.W., and Graybiel, A.M. (1993). Two input systems for body representations in the primate striatal matrix: experimental evidence in the squirrel monkey. J. Neurosci. 13, 1120–1137.

Flaherty, A.W., and Graybiel, A.M. (1995). Motor and somatosensory corticostriatal projection magnifications in the squirrel monkey. J. Neurophysiol. 74, 2638–2648.

Flinker, A., Korzeniewska, A., Shestyuk, A.Y., Franaszczuk, P.J., Dronkers, N.F., Knight, R.T., and Crone, N.E.(2015). Redefining the role of Broca’s area in speech. Proc. Natl. Acad. Sci. 112, 201414491.

Fogelson, N., Williams, D., Tijssen, M., van Bruggen, G., Speelman, H., and Brown, P. (2006). Different functional loops between cerebral cortex and the subthalmic area in Parkinson’s disease. Cereb. Cortex 16, 64–75.

Granger C.W.J. (1969). Investigating Causal Relations by Econometric Models and Cross-spectral Methods. Econometrica 37, 424–438.

Groppe, D.M., Urbach, T.P., and Kutas, M. (2011). Mass univariate analysis of event-related brain potentials/fields I: a critical tutorial review. Psychophysiology 48, 1711–1725.

Haynes, W.I.A., and Haber, S.N. (2013). The organization of prefrontal-subthalamic inputs in primates provides an anatomical substrate for both functional specificity and integration: implications for Basal Ganglia models and deep brain stimulation. J. Neurosci. 33, 4804–4814.

Herz, D.M., Siebner, H.R., Hulme, O.J., Florin, E., Christensen, M.S., and Timmermann, L. (2014). Levodopa reinstates connectivity from prefrontal to premotor cortex during externally paced movement in Parkinson’s disease. Neuroimage 90, 15–23.

Herz, D.M.M., Zavala, B.A.A., Bogacz, R., Brown, P., Herz, D.M.M., Zavala, B.A.A., Bogacz, R., and Brown, P.(2016). Neural Correlates of Decision Thresholds in the Human Subthalamic Nucleus. Curr. Biol. 26, 916–920.

Holm, S. (1979). A Simple Sequentially Rejective Multiple Test Procedure. Scand. J. Stat. 6, 65–70.

Jenkinson, N., Kühn, A.A., and Brown, P. (2013). γ oscillations in the human basal ganglia. Exp. Neurol. 245, 72–76.

Jensen, O., Goel, P., Kopell, N., Pohja, M., Hari, R., and Ermentrout, B. (2005). On the human sensorimotor-cortex beta rhythm: sources and modeling.Neuroimage 26, 347–355.

Kondylis, E.D., Randazzo, M.J., Alhourani, A., Lipski, W.J., Wozny, T.A., Pandya, Y., Ghuman, A.S., Turner, R.S., Crammond, D.J., and Richardson, R.M. (2016). Movement-related dynamics of cortical oscillations in Parkinson’s disease and essential tremor. Brain 139, 2211–2223.

Korzeniewska, A., Crainiceanu, C.M., Kuś, R., Franaszczuk, P.J., and Crone, N.E. (2008). Dynamics of event-related causality in brain electrical activity. Hum. Brain Mapp. 29, 1170–1192.

Korzeniewska A., Franaszczuk P.J., Crainiceanu C.M., Ku??, R., and Crone, N.E. (2011). Dynamics of large-scale cortical interactions at high gamma frequencies during word production: Event related causality (ERC) analysis of human electrocorticography (ECoG). Neuroimage 56, 2218–2237.

Lachaux, J.P., Rodriguez, E., Martinerie, J., and Varela, F.J. (1999). Measuring phase synchrony in brain signals. Hum. Brain Mapp. 8, 194–208.

Lalo, E., Thobois, S., Sharott, A., Polo, G., Mertens, P., Pogosyan, A., and Brown, P. (2008). Patterns of bidirectional communication between cortex and basal ganglia during movement in patients with Parkinson disease. J. Neurosci. 28, 3008–3016.

Lidsky, T.I., Manetto, C., and Schneider, J.S. (1985). A consideration of sensory factors involved in motor functions of the basal ganglia. Brain Res. 356, 133–146.

Lipski, W.J.W.J., Wozny, T.A.T.A., Alhourani, A., Kondylis, E.E.D., Turner, R.S.R.S., Crammond, D.J.D.J., and Richardson, R.M.R.M. (2017). Dynamics of human subthalamic neuron phase-locking to motor and sensory cortical oscillations during movement. J. Neurophysiol. 118.

Litvak, V., Jha, A., Eusebio, A., Oostenveld, R., Foltynie, T., Limousin, P., Zrinzo, L., Hariz, M.I., Friston, K., and Brown, P. (2011). Resting oscillatory cortico-subthalamic connectivity in patients with Parkinson’s disease. Brain 134, 359–374.

Litvak, V., Eusebio, A., Jha, A., Oostenveld, R., Barnes, G., Foltynie, T., Limousin, P., Zrinzo, L., Hariz, M.I., Friston, K.,et al. (2012). Movement-related changes in local and long-range synchronization in Parkinson’s disease revealed by simultaneous magnetoencephalography and intracranial recordings. J. Neurosci. 32, 10541–10553.

Manning, J.R., Jacobs, J., Fried, I., and Kahana, M.J. (2009). Broadband shifts in local field potential power spectra are correlated with single-neuron spiking in humans. J. Neurosci. 29, 13613–13620.

Maris, E., and Oostenveld, R. (2007). Nonparametric statistical testing of EEG-and MEG-data. J. Neurosci. Methods 164, 177–190.

Mazzoni, P., Hristova, A., and Krakauer, J.W. (2007). Why Don’t We Move Faster? Parkinson’s Disease, Movement Vigor, and Implicit Motivation. 27, 7105–7116.

Miller, K.J., Hermes, D., Honey, C.J., Hebb, A.O., Ramsey, N.F., Knight, R.T., Ojemann, J.G., and Fetz, E.E. (2012). Human motor cortical activity is selectively phase-entrained on underlying rhythms. PLoS Comput. Biol. 8, e1002655.

Montgomery, S.M., and Buzsáki, G. (2007). Gamma oscillations dynamically couple hippocampal CA3 and CA1 regions during memory task performance. Proc. Natl. Acad. Sci. U. S. A. 104, 14495–14500.

Nambu, A., Tokuno, H., Inase, M., and Takada, M. (1997). Corticosubthalamic input zones from forelimb representations of the dorsal and ventral divisions of the premotor cortex in the macaque monkey: comparison with the input zones from the primary motor cortex and the supplementary motor area. Neurosci. Lett. 239, 13–16.

Nishida, M., Korzeniewska, A., Crone, N.E., Toyoda, G., Nakai, Y., Ofen, N., Brown, E.C., and Asano, E. (2017). Brain network dynamics in the human articulatory loop. Clin. Neurophysiol. 128, 1473–1487.

Oswal, A., Beudel, M., Zrinzo, L., Limousin, P., Hariz, M., Foltynie, T., Litvak, V., and Brown, P. (2016). Deep brain stimulation modulates synchrony within spatially and spectrally distinct resting state networks in Parkinson’s disease. Brain 139, 1482–1496.

Panov, F., Levin, E., de Hemptinne, C., Swann, N.C., Qasim, S., Miocinovic, S., Ostrem, J.L., and Starr, P.A. (2016). Intraoperative electrocorticography for physiological research in movement disorders: principles and experience in 200 cases. J. Neurosurg. 1–10.

Parent, A., and Hazrati, L.N. (1995). Functional anatomy of the basal ganglia. II. The place of subthalamic nucleus and external pallidum in basal ganglia circuitry. Brain Res. Brain Res. Rev. 20, 128–154.

Randazzo M.J., Kondylis E.D., Alhourani A., Wozny T.A., Lipski W.J., Crammond D.J., and Richardson, R.M. (2016). Three-dimensional localization of cortical electrodes in deep brain stimulation surgery from intraoperative fluoroscopy. Neuroimage 125, 515–521.

Roberts, M.J., Lowet, E., Brunet, N.M., Ter Wal, M., Tiesinga, P., Fries, P., and De Weerd, P. (2013). Robust gamma coherence between macaque V1 and V2 by dynamic frequency matching. Neuron 78, 523–536.

Sherman, M.A., Lee, S., Law, R., Haegens, S., Thorn, C.A., Hämäläinen, M.S., Moore, C.I., and Jones, S.R.(2016). Neural mechanisms of transient neocortical beta rhythms: Converging evidence from humans, computational modeling, monkeys, and mice. Proc. Natl. Acad. Sci. 113, E4885–E4894.

Siegel, M., Donner, T.H., Oostenveld, R., Fries, P., and Engel, A.K. (2008). Neuronal Synchronization along the Dorsal Visual Pathway Reflects the Focus of Spatial Attention. Neuron 60, 709–719.

Smith, S.M., and Nichols, T.E. (2009). Threshold-free cluster enhancement: Addressing problems of smoothing, threshold dependence and localisation in cluster inference. Neuroimage 44, 83–98.

Tallon-Baudry, C., Kreiter, A., and Bertrand, O. (1999). Sustained and transient oscillatory responses in the gamma and beta bands in a visual short-term memory task in humans. Vis. Neurosci. 16, 449–459.

Tan, H., Pogosyan, A., Anzak, A., Ashkan, K., Bogdanovic, M., Green, A.L., Aziz, T., Foltynie, T., Limousin, P., Zrinzo, L.,et al. (2013). Complementary roles of different oscillatory activities in the subthalamic nucleus in coding motor effort in Parkinsonism. Exp. Neurol. 248, 187–195.

Tort, A.B.L., Kramer M. a, Thorn, C., Gibson, D.J., Kubota, Y., Graybiel, A.M., and Kopell, N.J. (2008). Dynamic cross-frequency couplings of local field potential oscillations in rat striatum and hippocampus during performance of a T-maze task. Proc. Natl. Acad. Sci. U. S. A. 105, 20517–20522.

Turner, R.S., and Desmurget, M. (2010). Basal ganglia contributions to motor control: a vigorous tutor. Curr. Opin. Neurobiol. 20, 704–716.

Voytek, B., Secundo, L., Bidet-Caulet, A., Scabini, D., Stiver, S.I., Gean, A.D., Manley, G.T., and Knight, R.T. (2010). Hemicraniectomy: a new model for human electrophysiology with high spatio-temporal resolution. J. Cogn. Neurosci. 22, 2491–2502.

Williams, D., Tijssen, M., Van Bruggen, G., Bosch, A., Insola, A., Di Lazzaro, V., Mazzone, P., Oliviero, A., Quartarone, A., Speelman, H.,et al. (2002). Dopamine-dependent changes in the functional connectivity between basal ganglia and cerebral cortex in humans. Brain 125, 1558–1569.

Witham, C.L., Wang, M., and Baker, S.N. (2010). Corticomuscular coherence between motor cortex, somatosensory areas and forearm muscles in the monkey. Front. Syst. Neurosci. 4.

Womelsdorf, T., Fries, P., Mitra, P.P., and Desimone, R. (2006). Gamma-band synchronization in visual cortex predicts speed of change detection. Nature 439, 733–736.

Yamawaki, N., Stanford, I.M., Hall, S.D., and Woodhall, G.L. (2008). Pharmacologically induced and stimulus evoked rhythmic neuronal oscillatory activity in the primary motor cortex in vitro. Neuroscience 151, 386–395.

